# Environmental fungi from cool and warm neighborhoods in the urban heat island of Baltimore City show differences in thermal susceptibility and pigmentation

**DOI:** 10.1101/2023.11.10.566554

**Authors:** Daniel F. Q. Smith, Madhura Kulkarni, Alexa Bencomo, Tasnim Syakirah Faiez, J. Marie Hardwick, Arturo Casadevall

## Abstract

A major barrier for most fungal species to infect humans is their inability to grow at body temperature (37°C). Global warming and more frequent extreme heat events may impose selection pressures that allow fungal adaptation to higher temperatures. Cities are heat islands that are up to 8°C warmer than their suburban counterparts because of mechanical heat production and reduced greenspace, among other factors, and may be an important reservoir of fungi that have increased risk of thermotolerance and inhabit environments near humans. Here we describe a novel and inexpensive technique that was developed to collect fungal samples from various sites in Baltimore, Maryland using commercially available taffy candy. Our results show fungal isolates from warmer neighborhoods show greater thermotolerance and lighter pigmentation relative to isolates of the same species from cooler neighborhoods, suggesting local adaptation. Lighter pigmentation in fungal isolates from warmer areas is consistent with known mechanisms of pigment regulation that modulate fungal cell temperature. The opportunistic pathogen *Rhodotorula mucilaginosa* from warmer neighborhoods had a higher resistance to gradual exposure to extreme heat than those from cooler neighborhoods. Our results imply fungal adaptation to increased temperature in an urban environment. The acquisition of thermotolerance poses a risk for humans if fungal species with pathogenic potential acquire the capacity to grow at human body temperatures and cause disease.

## Introduction

Humans endothermy provides natural protection against many fungal species with pathogenic potential [1]. Most fungi are unable to survive at or above human body temperature, and analyses of yeast thermal tolerance has shown that there is a 6% reduction in the number of fungal species able to survive at each degree above 30°C [1,2].

Natural and human-made disasters are associated with adaptive changes in environmental microbes that can lead to the emergence of new diseases in humans, plants, and animals [3]. Changes in the environment have been hypothesized to enable adaptive changes in fungi and drive the emergence of new fungal pathogens [4], a phenomenon that may have already occurred with *Candida auris* [4,5]. Global warming and extreme heat (including prolonged and more frequent heatwaves) may enable environmental fungi with pathogenic potential to acquire higher thermotolerance and thus overcome the mammalian thermal barrier. In recent decades, a sharp emergence of new fungal pathogens such as *Candida auris* and *Sporothrix brasiliensis* has been proposed to result from environmental pressure to acquire thermotolerance [4,6]. An analysis of the fungal collection at the Westerdijk Fungal Diversity Institute showed that fungal isolates collected globally from the environment in recent decades have a higher maximum growth temperature than those collected in prior decades, consistent with a possible gain of thermotolerance due to global warming [2]. However, that study is limited as the isolates in their collection are not randomly collected from the environment and may not represent the true diversity of fungi present in the environment over the course of a century.

Environments with high thermal selective pressure include urban centers (cities), which are subjected to the “heat island effect” and tend to be 4-8°C warmer than rural or suburban neighboring regions [7–10]. The warmer environment in cities is predominantly due to reduced green spaces, blocking of natural airflow by buildings, higher heat absorbance by building materials, and anthropomorphic heat production by machines and cars. However, even within cities, there can be large temperature variation, with some neighborhoods experiencing more severe heat than others, and the warmer neighborhoods are often the home to marginalized communities [10–12]. A study published by the National Oceanographic and Atmospheric Administration (NOAA) reported that in Baltimore City in late August 2018, parks and suburban neighborhoods are approximately 30.5°C, which is about 8-9°C cooler than the warmest neighborhoods that reach 39.4°C [13].

Currently, there are few studies investigating the mycobiome of urban centers, including areas where there are ample bacterial microbiome analyses, partly due to technical difficulties in performing metagenomic analysis of fungal DNA without adequate reference genomes for comparison and sequence alignment [14,15]. Few studies have investigated the heat-resistance and phenotypic profiles of urban fungi. An analysis of filamentous environmental fungi from urban sites in Louisville, Kentucky showed faster growth rates and higher enzymatic activities at higher temperatures than the fungi of the same species isolated from rural sites [16].

To understand mechanisms of the emergence of fungal pathogens there is a need for more functional characterization of urban mycobiome and a more detailed characterization of how heat affects urban fungal populations. Consequently, we explored the urban fungal census from sidewalks across thermally diverse neighborhoods in Baltimore, Maryland using a new technique for collection, and show examples of genotypic and phenotypic characterization of the fungi present, including pigmentation variation and the isolation and identification of clinically relevant polyextremophilic yeasts.

## Materials and Methods

### Sample Site and Date Selection

To select thermally diverse neighborhoods in Baltimore, MD, we used publicly available data on heat disparities across Baltimore City. Based on 2018 data from the National Oceanographic and Atmospheric Administration (NOAA) [13], we collected samples from four sites across Baltimore City. For the two warm neighborhood sites, we chose the corner of N. Wolfe St and E. Fayette St. (“Fayette St.”) and the corner of N. Charles St. and Washington Park East (“Mt. Vernon”), representative of the warmest areas in Baltimore. For the average temperature neighborhood site, we chose the corner of S. Wolfe St., and Fell St (“Fells Point”), and for a cool neighborhood site, we chose the corner of N. Charles St. and St. Paul St (“Guilford”). On August 20^th^, 2023, we collected samples from all four sites during sunny weather, between 12 P.M. and 2 P.M. Additional collections were done on September 4^th^, 2023, June 21^st^, 2024, and August 28^th^, 2024, between 12 P.M. and 3 P.M, for only the (warmer) Fayette St. and (cooler) Guilford sites. All dates were chosen due to sunny weather and were at least three days after the most recent precipitation. The two collection dates in June and August 2024 were for the purpose of performing further mold pigmentation measurements. Thermal images to record the temperature of each site were taken using a FLIR C2 IR camera, and images were processed using FIJI (ImageJ) IRimage plugin [17]. Additional temperature readings were done using a GP-300 infrared thermometer (Duraline Systems, West Nyack, NY) to confirm infrared camera measurements. Measurements for the thermometer and FLIR camera were comparable (difference <1°C) (Supplementary Table S1).

### Sidewalk Sample Collection

To collect fungal samples from sidewalks, we developed a collection protocol using Starburst taffy candy (Mars Corporation, VA, USA) (Figure 1A). Taffy is a soft, sticky candy that easily adapts its contour to that of environmental surfaces when pressed against those surfaces with hand pressure. The stickiness picks up small particles. To collect environmental fungi found in the gritty and textured surface of the sidewalk, we rolled a yellow Starburst taffy over an approximately 100 cm^2^ area on the sidewalk for 30 seconds, making sure to press the taffy into the pavement. The taffy candy was then dissolved in 10 mL PBS containing 60 µl of 25% KOH solution to neutralize the citric acid in the candy. After the taffy was dissolved, the suspension was passed through a 100 µm filter to remove debris, and 200 µl aliquots of sidewalk samples were spread on each of three to five 10 cm Sabouraud dextrose agar (SDA) plates with penicillin-streptomycin to limit bacterial growth. SDA was chosen as the isolation agar because it is acidic and inhibits many bacterial species while promoting fungal growth. Samples were incubated at room temperature (22°C) for up to 7 d. The Starburst taffy alone did not grow fungi when treated with these conditions, except for a single mold colony (Supplementary Figure 1).

**Figure 1.**
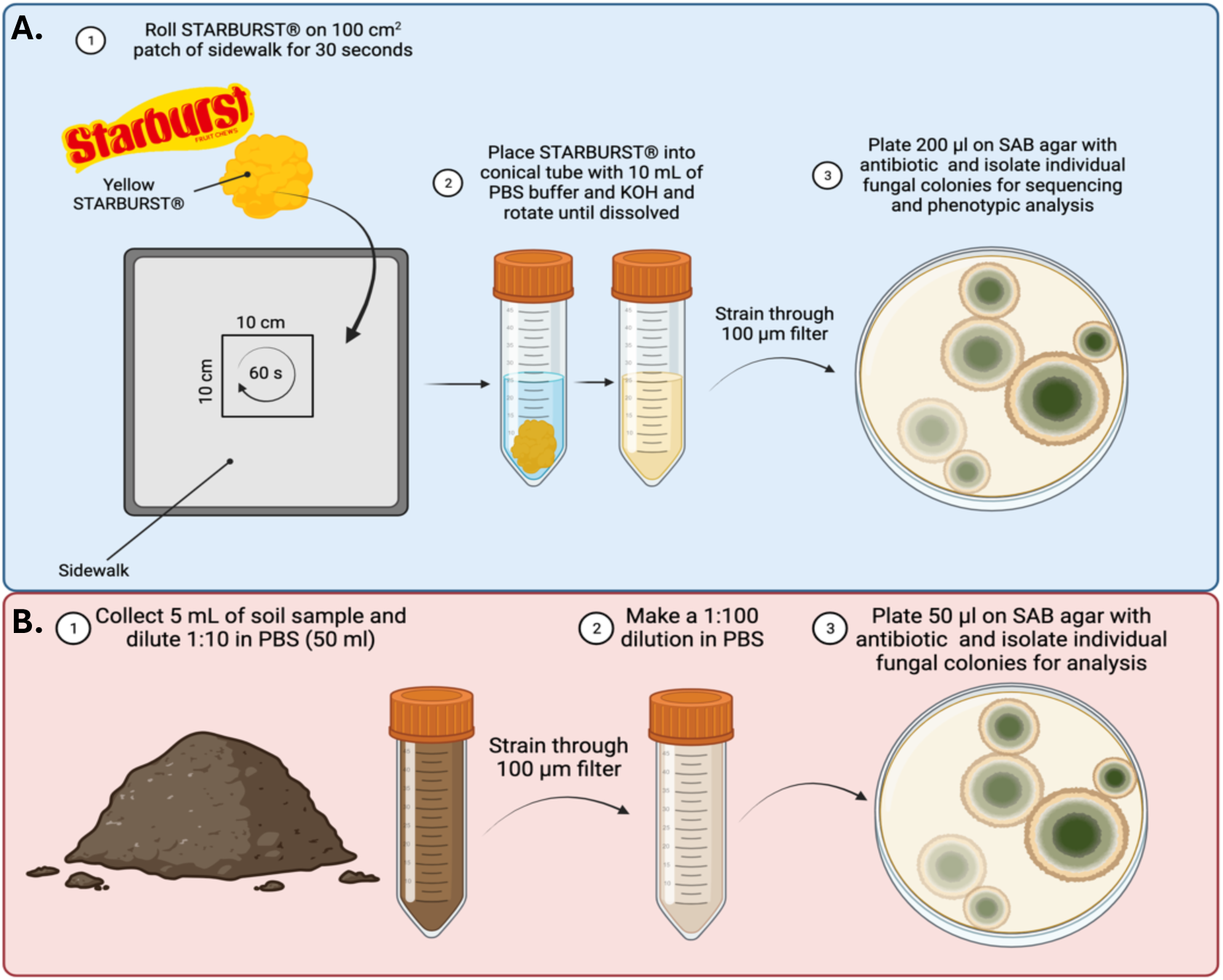
Sample Collection Methods. A. Fungal sample collection from the sidewalk via Starburst. B. Fungal sample collection from dirt.

### Dirt Sample Collection

Dirt samples were collected by scraping the top layer of dirt along the tree-wells, garden patches between sidewalks, and grass-tufted crevices adjacent to the curb. At least 5 mL of non-compacted dirt was collected at each location. The 5 mL of dirt sample were added to 45 mL PBS, solubilized at room temperature for 15-30 min, passed through a 100 µm strainer to remove large debris, plant material, and larger insoluble objects, diluted 1:100 in PBS, and 50 µl aliquots of diluted dirt samples were spread on 3-5 10 cm SDA plates containing penicillin-streptomycin and incubated at room temperature (22°C) for up to 7 d. Isolation scheme summarized in Figure 1B.

### Yeast Colony Isolation

Fungal colonies were isolated from original SDA plates containing material from sidewalk and dirt samples by re-streaking on SDA plates. Plates were incubated at room temperature for up to 7 d. Plates were examined daily for fungal growth, both filamentous and yeast like. Yeast-like colonies (small, circular, opaque, and non-filamentous) were picked with a pipette tip and streaked to isolation on fresh yeast peptone dextrose (YPD) agar plates, and incubated at 20°C. Newly appearing colonies derived from the dirt and sidewalk collection plates were picked from the YPD plates daily for up to 7 d. Cells from each collected isolate were observed microscopically to confirm fungal (non-bacterial) identify by cell size, morphology, and appearance of intracellular organelles. Individual fungal colonies were isolated as they appeared, transferred to 96-well plates with freezing media (25% glycerol, 75% YPD broth), and stored at −80°C. Replicate 96-well plates with sterile PBS were inoculated and stored at 4°C. All original plates were wrapped in parafilm and stored at 4°C.

### Growth Area Measurements

All the original samples on SDA agar plates from Fayette St. and Guilford were scanned using a Canon CanoScan9000F scanner at 600 dpi after 7 days of growth. Using the FIJI (ImageJ) selection and measure tools, ROIs were manually identified and the total area of fungal growth per plate was determined for each site.

### Pigmentation Analysis

Pigmentation of fungi was determined as previously described [18]. Briefly, pigmentation of molds growing on the original SDA sample plates was assessed by imaging with a Canon CanoScan9000F scanner at 600 dpi. To measure the pigmentation of yeast isolates, frozen stocks of isolated colonies stored at −80°C were stamped onto YPD agar on an OmniWell agar plate using a flame-sterilized microplate replicator (Boekel Scientific) and grown at room temperature for 7 days. Yeast growth (which were in spots of ∼ 5 mm in diameter) were scanned using the CanoScan9000F at 600 dpi. Images were processed and mean gray values of mold and yeast colonies were measured using FIJI (ImageJ) measure tool.

### Thermal Absorbance Analysis

Agar plates with mold cultures were acclimated to room temperature for 1 h (22°C). Petri dishes were unlidded and exposed to a pre-warmed 19W 4000K white light bulb at 50 cm for 10 minutes. Following the 10 min exposure, culture plates were imaged immediately for heat absorption using an FLIR E96 infrared camera. Images were processed using FIJI (ImageJ) IRimage plugin [17], and temperature measurements were recorded for each mold colony. Mean gray value measurements were taken using the visible light spectrum image taken by the FLIR E96 camera at time of experiment.

### Heat-Ramp Assay

Heat-Ramp experiments were performed as previously described, with some changes [19]. Frozen stocks of *R. mucilaginosa* and *C. minutum* collected from Fayette St., Mt. Vernon, and Guilford were streaked on SDA plates and incubated 48 h at room temperature. Single colonies of *R. mucilaginosa* and *C. minutum* were scraped from SDA plates, suspended in liquid SDA and adjusted to a density of 0.1 absorbance at 600 nm and 200 µl of these cultures were grown overnight at 30°C in stationary 96-well plates. Samples were diluted 1:5 in fresh SDA liquid medium, and 100 µl of these cultures were directly treated with a linear heat-ramp from 30°C to 55, 56, or 60°C over 8-10 min in a water bath with agitation (LAUDA Scientific, Germany). Untreated and heat-ramp–treated strains were immediately spotted (5 μL in SDA) in five-fold serial dilutions on SDA agar and incubated at 30 °C for 48 h. Plates were imaged and CFUs were counted for each isolate under each treatment.

### Internal Transcribed Spacer (ITS) Sequencing

Colonies for each isolate were picked from newly streaked SDA plates of fungal isolates touched lightly with a pipette tip and transferred and deposited along the inside of an empty PCR tube in a thin layer. For cultures that were unable to grow following storage at −80°C, we scraped some of the frozen stock and deposited it inside of a PCR tube. The tubes were microwaved on high (1100 W) for 2 minutes as previously described [20]. DreamTaq Green PCR Master Mix (ThermoFisher) was added to each tube with 0.2 µM ITS4 and ITS5 primers to detect species-specific fungal internal transcribed sequences. The sequences of these primers are as follows: ITS4 - 5′-TCCTCCGCTTATTGATATGC-3′ and ITS5 - 5′-GGAAGTAAAAGTCGTAACAAGG-3′. PCR products were amplified according to the manufacturer’s protocol; 3 min initial denaturation at 95°C, 35 cycles with 30 s denaturation at 95°C, 30 s annealing at 55°C, and 1 min extension at 78°C, with a final 5 min extension at 78°C. PCR products were run on 1% agarose gel to confirm product amplification, and Sanger sequenced at the Johns Hopkins School of Medicine Sequencing & Synthesis Core facility. Resulting sequences were analyzed using 4Peaks software and NCBI Blast nucleotide feature.

## Results

To investigate the potential effects of urban heat islands on fungi, we collected samples from four sites (Figure 2A and 2B) across Baltimore City based on 2018 air temperatures from the National Oceanographic and Atmospheric Administration (NOAA) [13]. Fungal samples were collected between 12 and 3 P.M. in Baltimore City on four dates with reported air temperatures of 27-37°C during the summers of 2023 and 2024. Sample sites represented both warmer and cooler neighborhoods and each site was representative of the environment in its neighborhood based on NOAA heat island maps. The four sites chosen for the study were: Site 1, (Fells Point) representing an average air temperature neighborhood, Site 2 (Fayette St.) representing the warmest area within Baltimore City, Site 3 (Mt. Vernon) representing an area with above average temperature, and Site 4 (Guilford) representing a neighborhood with cooler temperatures.

**Figure 2.**
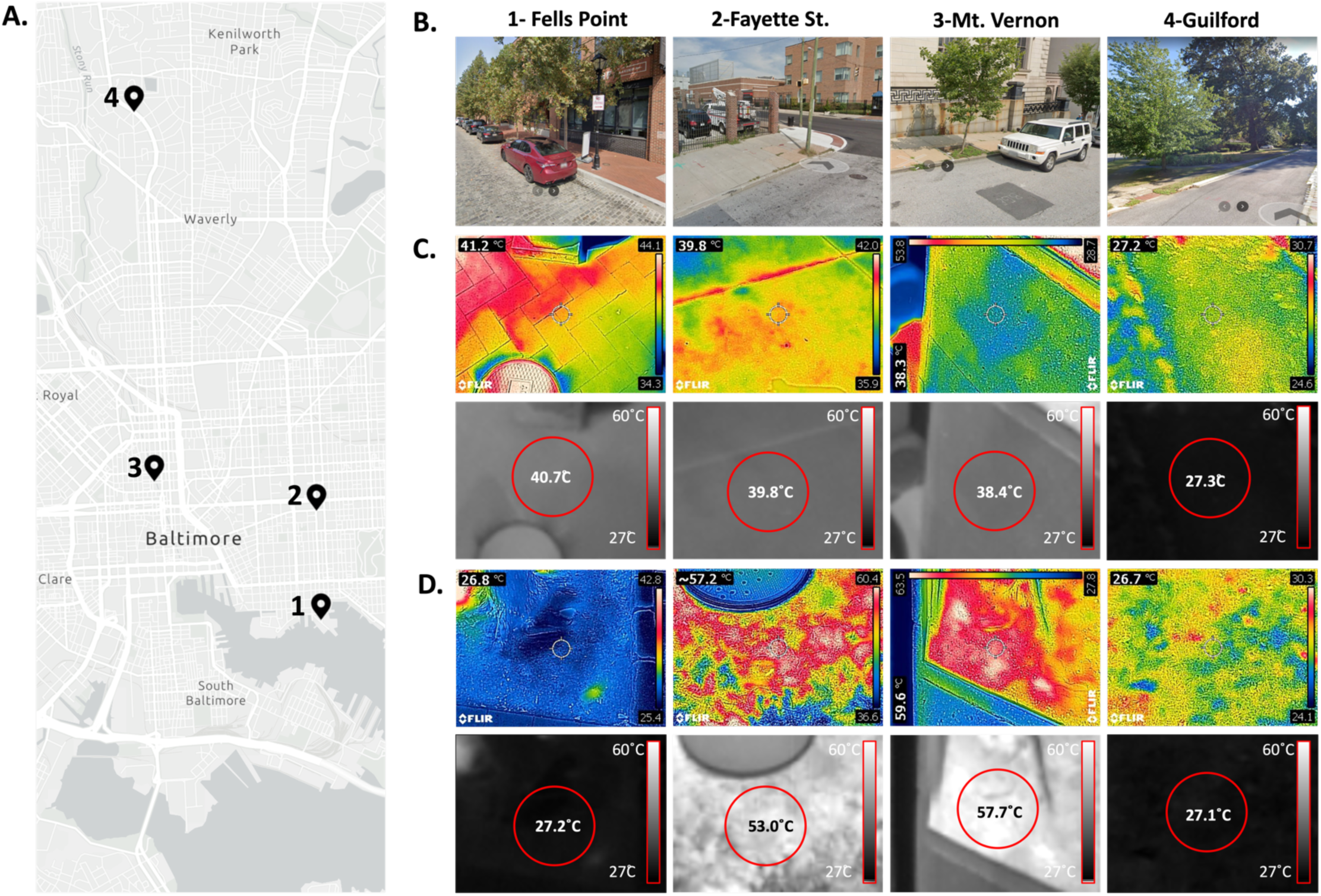
Thermal Properties of Sampling Sites in Baltimore, MD. A. Map of the four sites sampled in Baltimore, MD (Fells Point, Fayette, Mt. Vernon, Guilford). B. Google StreetView images from the sampling sites taken during summer months in recent years. Infrared images of the sidewalks (C) and dirt (D) on August 20^th^, 2023. The visible light photo from the FLIR 96E camera overlayed on the generated thermal image (top rows of C and D), and the raw grayscale temperature image standardized to 27°C (min) and 60°C (max) with the mean temperature from the center (approximate area shown with a red circle) of the individual image measured overlayed (bottom rows of C and D).

At each collection site, we measured sidewalk and dirt surface temperatures (Supplementary Table 1). The surface temperatures we observed were consistent with patterns we expected based on heat-island effect air temperatures published by NOAA. On the first and second collection days (August 20, 2023, and September 4^th^, 2023), we collected temperature data using an FLIR E96 camera to verify the temperature patterns. On the second through fourth collection dates, surface temperatures were recorded using a GP-300 infrared thermometer. The raw images from the infrared camera to measure surface temperatures on August 20^th^, 2023 are displayed in the upper panels of Figure 2C and D, with an automatically scaled color range based on the temperature range in the field of view, which differs in each frame. To more directly compare average temperatures between the dirt and sidewalk sample collection sites, the raw image files were converted to the gray scale (Figure 2C and 2D, lower rows) to reflect the absolute temperature range. The grayscale image allows for direct comparisons and quantifications, as it is scaled so that the gray value images have 27°C as the darkest pixel value and the whitest pixel values to represent 60°C. The measured average temperature from the approximate area of the red circle at the center of the image is overlayed.

On August 20^th^, 2023, the Fells Point site had a warm sidewalk with an average of 40.7°C, while a cooler dirt sample ambient temperature (27.2 °C) (Figure 2C, D). As expected, the dirt and sidewalks from the Fayette St. and Mt. Vernon collection sites had the warmest surfaces tested. Sidewalks in these locations were ∼38.4-39.8°C (Figure 2C), with the dirt temperature at either site of sample collection reaching up to ∼53-57.7°C on average, with some areas on the surface reaching up to 60°C (Figure 2D). Both sidewalk and dirt samples from Guilford were ∼27°C, which was approximately the same as the reported air temperature at the time of collection (Supplementary Table 1), and shows a reduced thermal pressure compared to the high surface temperatures at other areas within the city.

In the subsequent sample collections on September 4^th^, 2023, June 21^st^, 2024, and August 28^th^, 2024, we collected only from Fayette St. and Guilford sites to simplify the collection and comparison process on the basis that they are on two ends of the heat-island landscape within Baltimore City, with Fayette St. being amongst the warmest areas and Guilford being amongst the coolest areas. In the subsequent fungal sample collection dates at the Fayette St. and Guilford sites, surface temperatures recorded via infrared thermometers remained consistent with what we observed on August 20^th^, 2023, and September 4^th^, 2023, using the infrared camera (Supplementary Table 1).

We successfully isolated yeast and mold from all four sites on August 20^th^, 2023, and from Guilford and Fayette sites on the remaining collection dates, including the sidewalk and dirt that reached exceedingly high surface temperatures. While fungi were isolated from samples across all neighborhoods, the abundance of culturable fungi varied by location. Notably, for the samples plated from sidewalk and dirt from four collection dates, less area of the petri dish cultures was covered in mold and yeast in plates plated with material from the warmest (Fayette St.) samples than the plates from the coolest (Guilford) site (Figure 3). While we were able to recover viable fungi at these extreme temperature environments in cities, there was a reduced amount of viable and culturable fungi present in the hotter areas. This method indicates that there are fewer culturable fungi present in the warmer neighborhoods, but this method does not account for the presence of species that may have a faster growth rate or species that may have a naturally larger growth area, causing increased relative coverage of the media plate.

**Figure 3.**
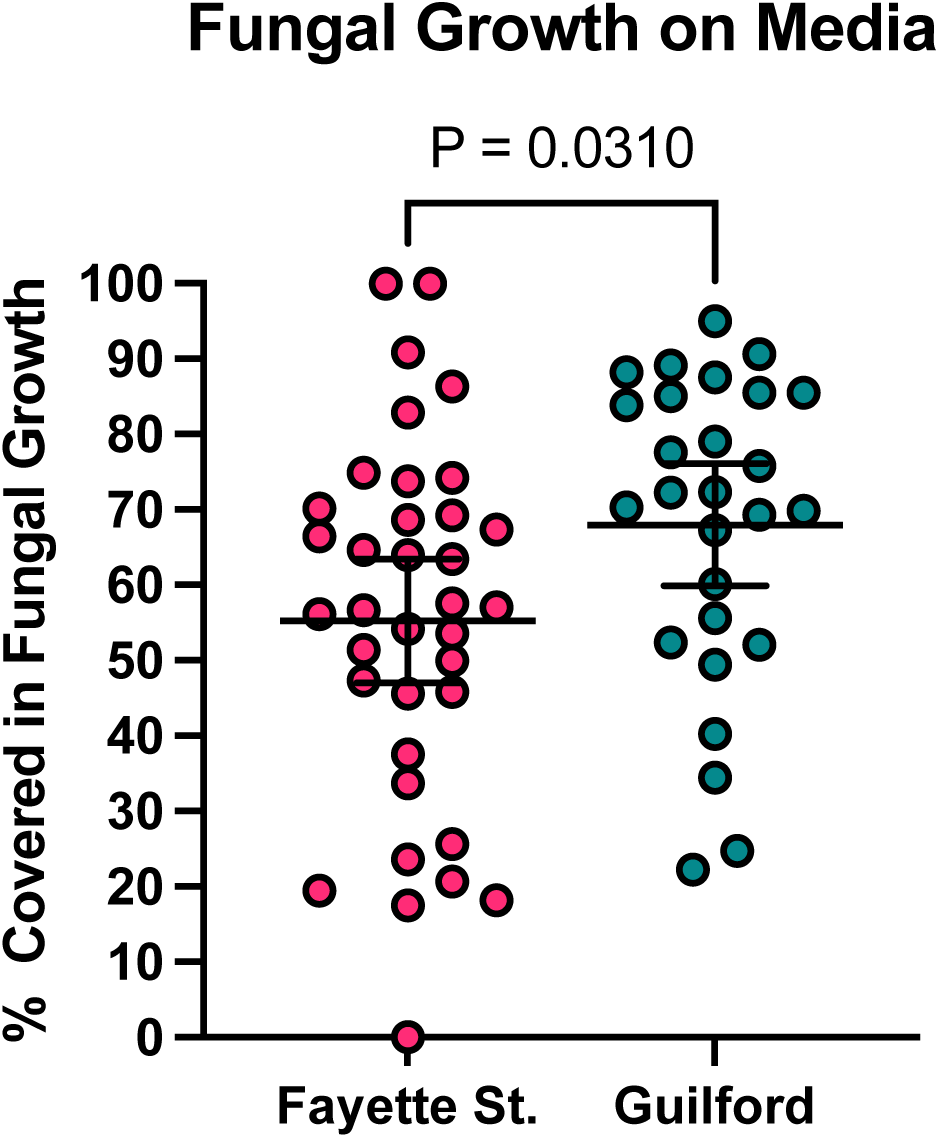
Fungal growth is sparser from Fayette St. (warm) Samples than Guilford (cool) Samples. Less plate surface area is covered by fungal growth from samples collected from Fayette St. compared to those collected at Guilford. Error bars represent 95% CI from the mean. Samples were collected on four collection dates. Each dot represents the percentage of agar area covered in fungal growth from one plate. Statistical significance was calculated with an unpaired t-test.

Pigments produced by fungi can absorb and retain heat, which impacts the cell temperature and supports fungal growth in a heat-deprived environment through a process described as “thermal melanism” [21]. Dark pigments in fungi are often composed of melanins, which have a remarkable capacity to absorb electromagnetic radiation and convert it to heat [18,21]. We compared mold pigmentation across the urban landscape from sidewalk and dirt on all 4 collection dates and found that molds from the Fayette St. site (warm – 40°C-57°C) were significantly less pigmented (lighter in color) than those from the Guilford site (cool – 27°C-33°C), with mean gray values differing by ∼10-pixel values (Figure 4A). Similarly, yeast from the first two collection dates were about 12-pixel values lighter from Fayette St. (warm) compared to Guilford site (cool) (Figures 4A and B). As an alternative approach, we directly assessed fungal heat absorbance by mold colonies as measured by an infrared camera. Consistent with having darker pigmentation, the mold found in the Guilford site (cool) absorbed more heat during a 10-minute exposure to white light than the mold from Fayette St. (warm) (Figures 4C and D). We would expect similar effects for yeast, though their small size precluded analyses by this method. Three mold plates each from samples collected from Fayette St. and Guilford collected on August 20^th^, 2023, and September 4^th^, 2023, were chosen at random. There was a correlation between the thermal absorbance (temperature) following exposure to white light (Figure 4C) and the mean gray value of the colonies (Figure 4E). This correlation is suggestive of thermal melanism/thermal albinism, as these phenomena have been called, to regulate thermal properties of the fungi [18,21]. These pigmentation patterns have been previously described for fungi collected at different latitudes, where fungi from arctic regions display enhanced pigmentation that promotes heat absorbance, while equatorial fungi have less pigmentation to avoid excessive heat absorbance [18].

**Figure 4.**
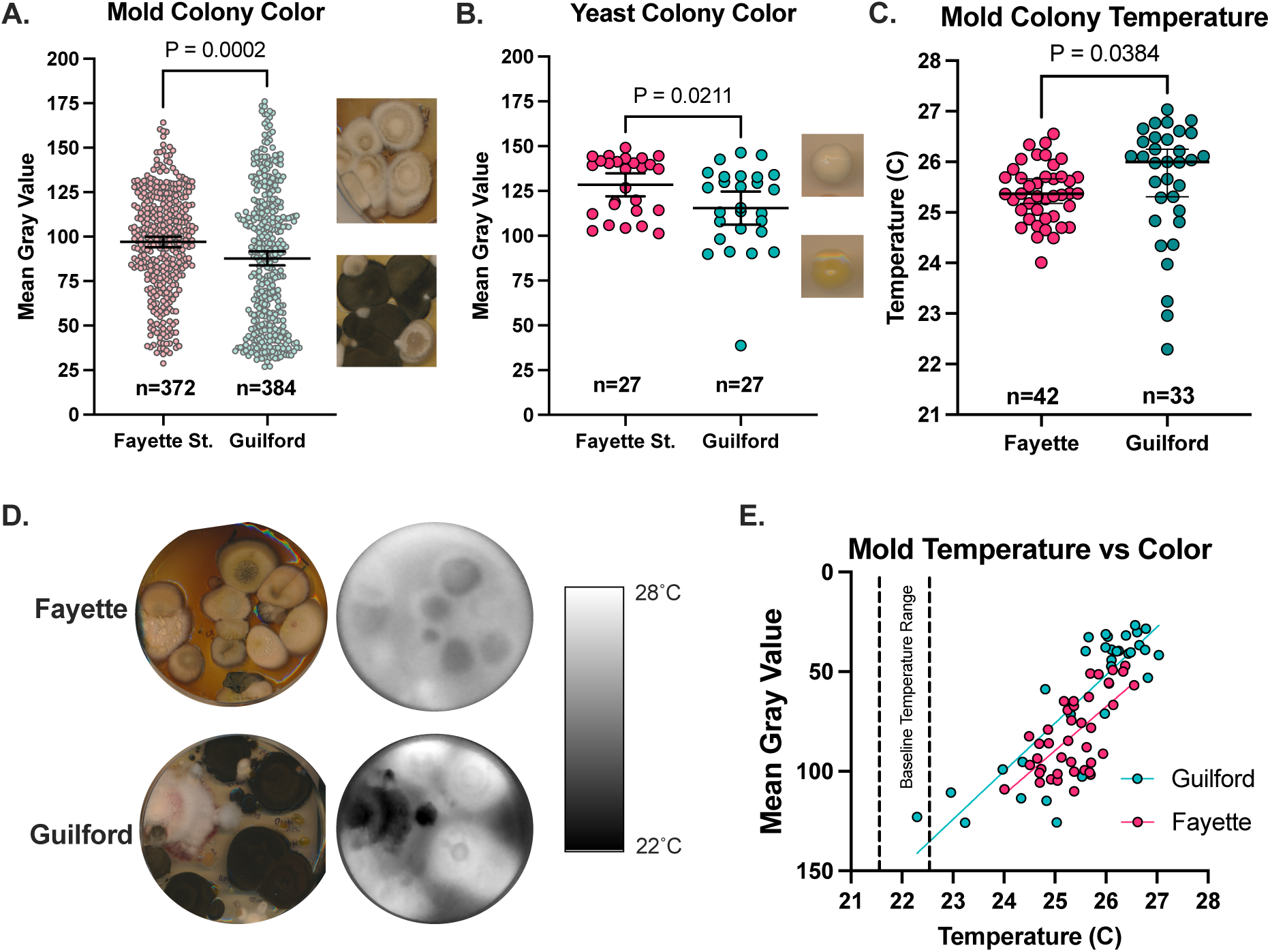
Fungi collected in thermally diverse neighborhoods exhibit thermal and pigment differences. The mean gray value of mold (A) and yeast (B) isolated from Fayette St. is higher than the mean gray value of the fungi isolated from Guilford, indicating less pigmentation in the Fayette St. fungi. Each dot represents values from individual colonies. Following exposure to light, fungi from the Guilford (cool) samples get warmer than those from the Fayette samples (Panels C and D). Correlation between colony temperature and the mean gray value, where the darker colonies get warmer (Panel E). Panels A and B show mean values with 95% CI and t-tests, while C shows median temperature and non-parametric Mann-Whitney test. Each dot represents mean gray value (Panel A, B, E) or temperature value (Panel C, E) for an individual mold (Panel A, C, E) or yeast colony (Panel B). Samples in (A) were collected from August 20^th^, 2023, September 4th, 2023, June 21st, 2024, and August 28th, 2024.Samples in (Panels B to E) were collected on August 20^th^, 2023, and September 4th, 2023.

To further investigate and compare the thermal properties of the fungi collected from warmer or cooler sites, we looked for culturable yeast of the same or closely related species by PCR amplifying and sequencing the ITS sequence of the 107 collected fungal isolates. We identified the yeast isolates by the species that most closely matched the Internal Transcribed Spacer (ITS) DNA sequence on NCBI GenBank, usually with high sequence identity. Notably, the presence of *Cystofilobasidium macerans, Rhodotorula spp., Cystobasidium spp.,* and other fungi in the Cystobasidiomycetes class was found in all neighborhood sampling sites (Supplementary Table 2). This class of yeast is ubiquitous in the environment, and some species are rare human pathogens. The identification of the same fungal species in different sampling sites allowed us to compare the characteristics of the same species of fungi when exposed to different environmental pressures. We found *R. mucilaginosa,* a ubiquitous environmental yeast and rare human pathogen, in both Fayette St. (warm) and Guilford (cool). We tested whether the *R. mucilaginosa* isolate from the Fayette St. location (Isolate 200S) had a different susceptibility to heat stress compared to those isolated from Guilford (414S, 423S, and 432S) in laboratory settings. We found that while none were able to grow at 37°C, the Fayette St. 200S isolate had greater resistance to killing following exposure to a 30-55°C and 30-56°C heat-ramp stress, remaining more viable compared to the three isolates from Guilford (Figure 5A, B).

**Figure 5.**
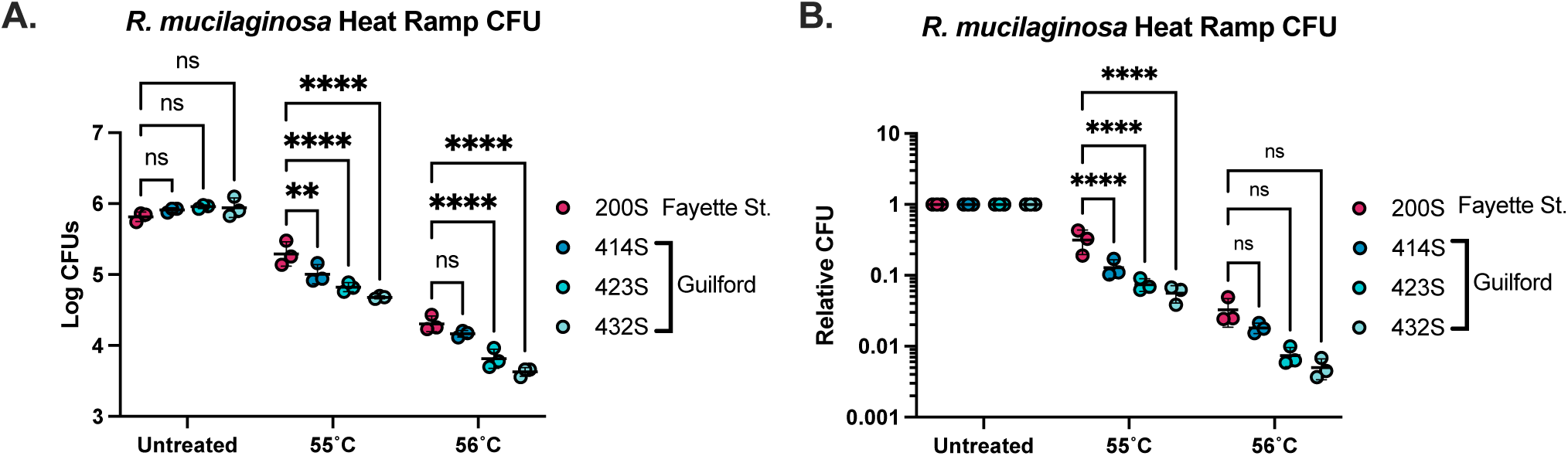
*R. mucilaginosa* from warmer neighborhoods has greater resistance to heat ramp stress. *R. mucilaginosa* from the sidewalk of Fayette St. (200S) show increased resistance to heat-ramp cell death stress compared to *R. mucilaginosa* strains collected from the cooler Guilford sidewalk (414S, 423S, and 432S) after gradual heat ramp to 55°C (Panels A and B). Significance of Two-Way ANOVA with multiple comparisons to the 200S for A and B at each condition with three biological replicates. Each dot represent CFUs from an individual biological replicate. **** represents p<0.0001, ** represents p<0.01, and ns represents p >0.05.

We found fungi belonging to the *Cystobasidium* genus, which includes *C. minutum,* a rare opportunistic fungal pathogen [22]. These fungi were found at was found at the two warmest sites, Fayette St and Mt. Vernon. One of the isolates of *C. minutum* recovered from the

38.4°C sidewalk average temperature in Mt. Vernon (300S) was thermotolerant (able to grow at 37°C), while another *C. minutum* isolate from Mt. Vernon sidewalk (305S) and a *C. lysinophilum* isolate from Fayette St. dirt (242D) were unable to grow at the higher temperatures (Supplementary Figure 2A). We performed similar experiments with *C. minutum* isolated from Mt. Vernon sidewalk (300S and 305S) and found that the 300S Mt. Vernon isolate was more resistant to heat-ramp at 55°C and 60°C (Supplementary Figure 2B,C). In fact, the 300S *C. minutum* isolate was the only one able to grow after brief exposure to 30-60°C heat-ramp. Together, these data suggest that even within one geographic location, a subset of yeast belonging to the same species are undergoing adaptation or selection for extreme heat.

Of the 44 yeast-like isolates collected from Fayette St. in our study on August 20^th^, 2023 and September 4^th^, 2023, 13 (29.5%) were identified as *Aureobasidium pullulans*. The identity of this fungus was established using ITS sequencing, showing conserved ITS sequence to the closest sequence match *A. pullulans* PE_11 strain (Supplementary Table 2, Supplementary Figure 3). *A. pullulans* is notable as it is a polyextremotolerant fungus, including temperature, saline stress, and nutrient stress, having first been isolated in glacial ice [23]. Microscopically, we observed *A. pullulans* to have diverse morphology with hyphae, yeast-like structures, and other abnormal shapes (Supplementary Figure 3). Additionally, 5 of the 44 isolates (11%) were identified as *Zalaria obscura,* a fungus that is closely related to *A. pullulans* and has similar resistances to environmental stressors [24]. Conversely, only 2 out of the 51 (4%) of the isolates from Guilford were identified as *A. pullulans (*Supplementary Table 2). Identification of these species in a neighborhood experiencing the most extreme heat in Baltimore City is thus a notable finding and places further context in this fungus’ role as a potential human pathogen.

## Discussion

As areas vulnerable to extreme heat conditions, cities are potential crucibles for the adaptation of fungi to warmer temperatures, and this can include fungal species with pathogenic potential that could lead to the emergence of new human fungal pathogens. Here we investigated the thermal tolerance of fungi in various neighborhoods of a city using a new to collect and culture environmental fungi. We found that fungi from warmer neighborhoods manifested different thermal tolerances and pigmentation from those of colder neighborhoods and our study suggests that the extreme ranges in heat in neighborhoods within urban centers may be sufficient to drive similar differences. The temperatures of the sidewalk and dirt in the warmer Fayette St. location were 40°C and 55°C, respectively, which are both above the typical temperature ranges of fungi, and above human body temperature (37°C), thus providing a selective pressure for higher heat tolerances that, if it selects for higher thermotolerance, could allow them to survive at mammalian body temperature. Meanwhile, the temperatures at the cooler, more wooded and shaded, Guilford neighborhood were around 27°C, and thus did not provide a selective pressure to for growth at or above mammalian body temperature. We noted lower fungal mass growing in agar from warmer sites, consistent with a reduced mycotic flora at those sites. Given that many fungal species are naturally hypothermic [25], and that the fungal viability of fungal species declines rapidly about 30°C [1], one might anticipate that the reduced fungal census in warmer sites reflects a reduced ability to handle heat stress.

*R. mucilaginosa* isolated from Fayette St (warm) was better equipped to survive gradual exposure to extreme heat (55°C) compared to isolates of the same species isolated from the Guilford site (cool). This suggests adaptation to the greater temperatures of the sidewalk at Fayette, implying selection for *R. mucilaginosa* that can withstand extreme thermal environments for short periods of time although all four were unable to grow at 37°C. This potential adaptation in *R. mucilaginosa* to heat is of interest since this fungus causes rare cases of systemic infection in immunocompromised individuals [26–28], and thus demonstrates some ability to withstand growth at human body temperature and host immune defenses. The addition of enhanced thermotolerant properties due to extreme environmental heat may allow this fungus to be even more equipped to survive within mammalian hosts. A relative of *Rhodotorula, Rhodosporidiobolus fluvialis,* was recently found in two independent clinical infections in China [29]. Researchers found that exposure of this fungus to 37°C conditions induced genetic and phenotypic changes that caused the fungus to have enhanced thermotolerance, drug resistance, and higher virulence phenotypes. These findings support the idea that fungi related to *Rhodotorula spp.,* are in a position to adapt to a warming environment, which can lead to enhanced frequency or severity of cases of disease with this fungus. *Rhodotorula spp.,* is found in nearly every environmental niche across the globe, and is particularly enriched within the guano of the lesser black-backed gull *Larus fuscus* [30]. This finding is interesting for several reasons. Firstly, the guano samples were specifically collected from urban sites suggesting that gulls may disemminate *Rhodotorula* within cities throught their guano, as was with pigeons and *Cryptococcus neoformans* [31]. Secondly, the internal temperature of *L. fuscus* is 41.2°C [32], which is over human body temperature. This indicates that *Rhodotorula* may have a natural ability to withstand, if not grow, at higher temperatures within an organism, and be primed for inhabiting warm urban sidewalks upon extrection.

We found a strong presence of the extremotolerant fungus *A. pullulans* in hotter neighborhoods. This fungus is capable of surviving saline, thermal, and nutrient-poor stresses, and can also be an opportunistic human pathogen [33–35]. The presence of this fungus is thus notable as it indicates the selection for extremotolerant fungal populations, which may correlate to ability to tolerate other stressors including mammalian immune response. Gostinčar et. al. has noted the potential importance of *A. pullulans* as an emerging pathogen due to its ability to resist extreme conditions, hyperplasticity, and its presence within the indoor built environment [23].

The previous comparative study of urban and rural fungi [16], along with the longitudinal analysis of thermal tolerance in culture collection isolates [2] and our results in this study, are each consistent with the notion that fungi are adaptingto our changing climate. We have found evidence that urban fungi on the sidewalk and soil of warmer neighborhoods have adaptations that allow them to mitigate heat and survive exposure to extreme heat. Molds and yeast in warmer neighborhoods had reduced pigmentation to avoid absorbing heat, while species like *R. mucilaginosa,* survive exposure to extreme temperatures (55°C) better than those from cooler sites. Even within warmer sites, fungi like *C. minutum* appear to show varied adaptation to heat where some isolates can survive up to a 60°C heat-ramp while another cannot, even though they are presumably recently related. Within these environments, we also see polyextremotolerant fungi such as *A. pullulans.* Many of these neighborhoods experiencing extreme heat vulnerability (and the risk of emerging thermotolerance in fungi), are also often inhabited by marginalized communities [10,12,36]. This is particularly worrisome, as these communities are also the ones already most vulnerable to fungal infection due to healthcare disparities and other social determinants of health [37].

In summary, our study shows differences in thermal tolerance for fungal species within a city depending on the neighborhoods of origin and their average ambient temperatures. Our findings warn that the heat islands defined by cities could be sites for rapid fungal adaptation to higher temperatures, which brings the possibility that some fungal species with pathogenic potential may be able to overcome human thermal defenses. Given that humans and fungi in city ecosystems are in close proximity to one another and that the number of immunocompromised patients are increasing yearly [38], rapid fungal thermal evolution in city heat islands has the potential to bring us new fungal diseases in the years ahead.

## Acknowledgements

We would like to acknowledge C. Johnson, Gracen Gerbig, and Dr. Jenna Glatzer for the assistance and support in collecting the fungal samples, Dr. Radames Cordero for the guidance related to thermal melanism in the fungal isolates. Figure 1 was made using BioRender.com. A.C. and D.F.Q.S. were supported in part by NIH grants AI162381, AI152078, and HL059842, AI168539 and AI183596. Funders played no role in the experimental design or outcome of the project.

---

**Supplementary Figure 1.**
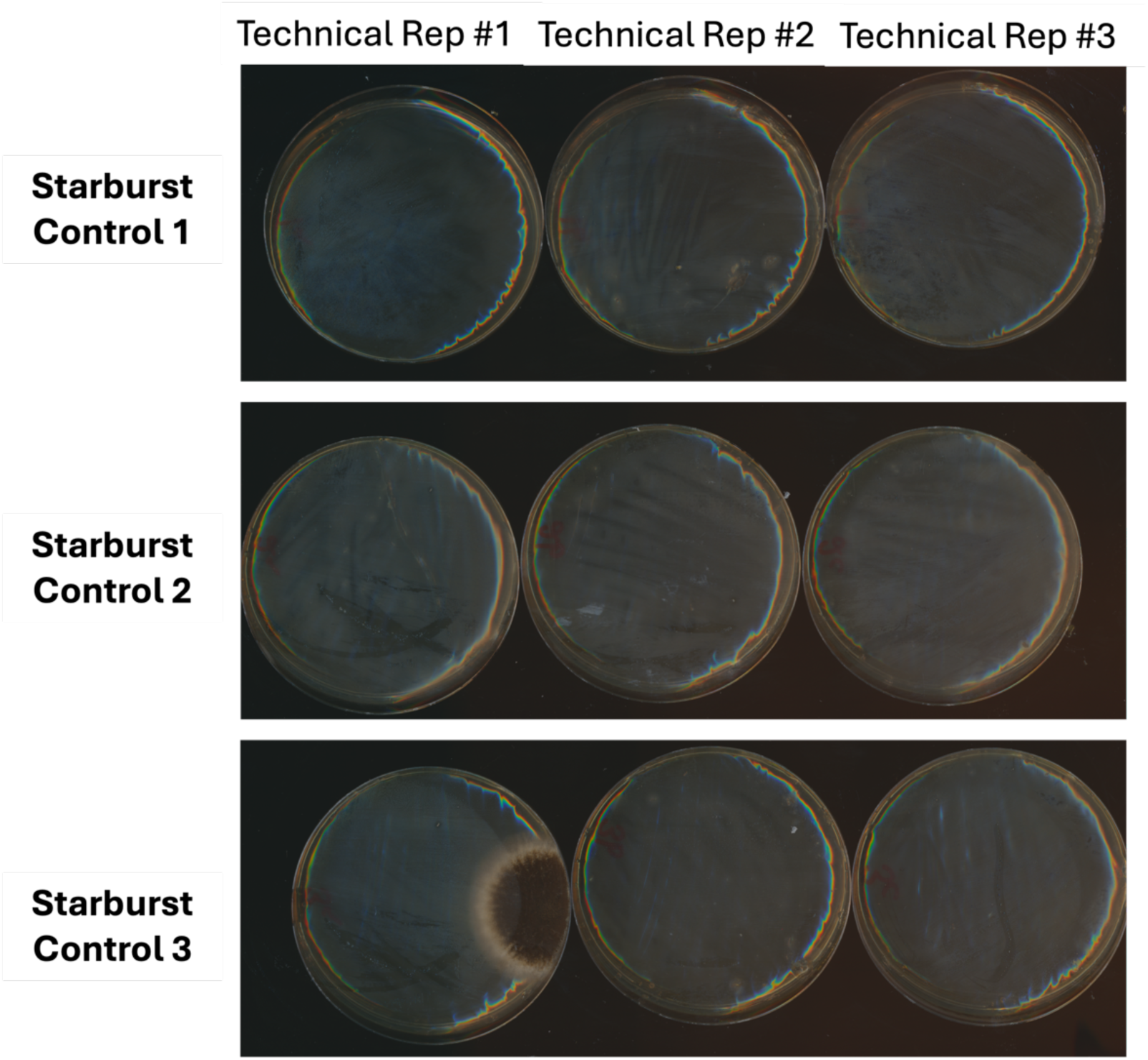

**Supplementary Figure 2.**
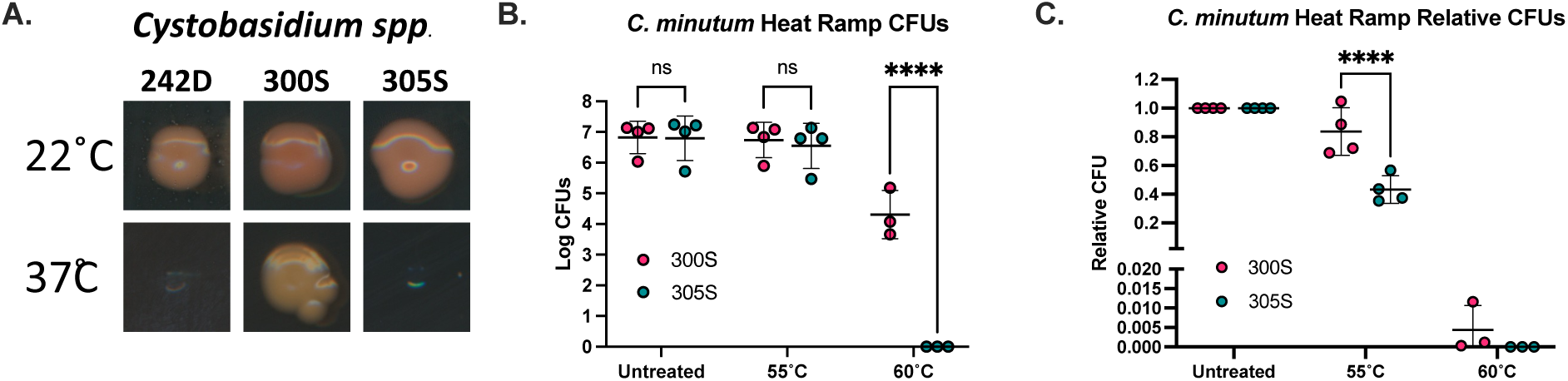
Thermotolerance of Cystobasidium spp. **(A)** Cystobasidium spp. isolated from Fayette St. dirt (242D) and Guilford sidewalks (*305*S) were unable to grow at 37°C, but *one* C. minutum isolate from Mt. Vernon sidewalk (30*0*S) *was* thermotolerant at 37°C. *C. minutum* from the sidewalk of Mt. Vernon (300S) grows better than the other C. minutum isolated from the Mt. Vernon sidewalk (305S) when exposed to gradual 55°C heat ramp, while only 300S can grow following exposure to 60°C heat ramp (Panels B and C). *S*ignificance of Two-Way ANOVA with multiple comparisons to the 300S for B and C at each condition with three biological replicates. Each dot represent CFUs from an individual biological replicate. **** represents p<0.0001, ** represents p<0.01, and ns represents p >0.05.

**Supplementary Figure 3.**
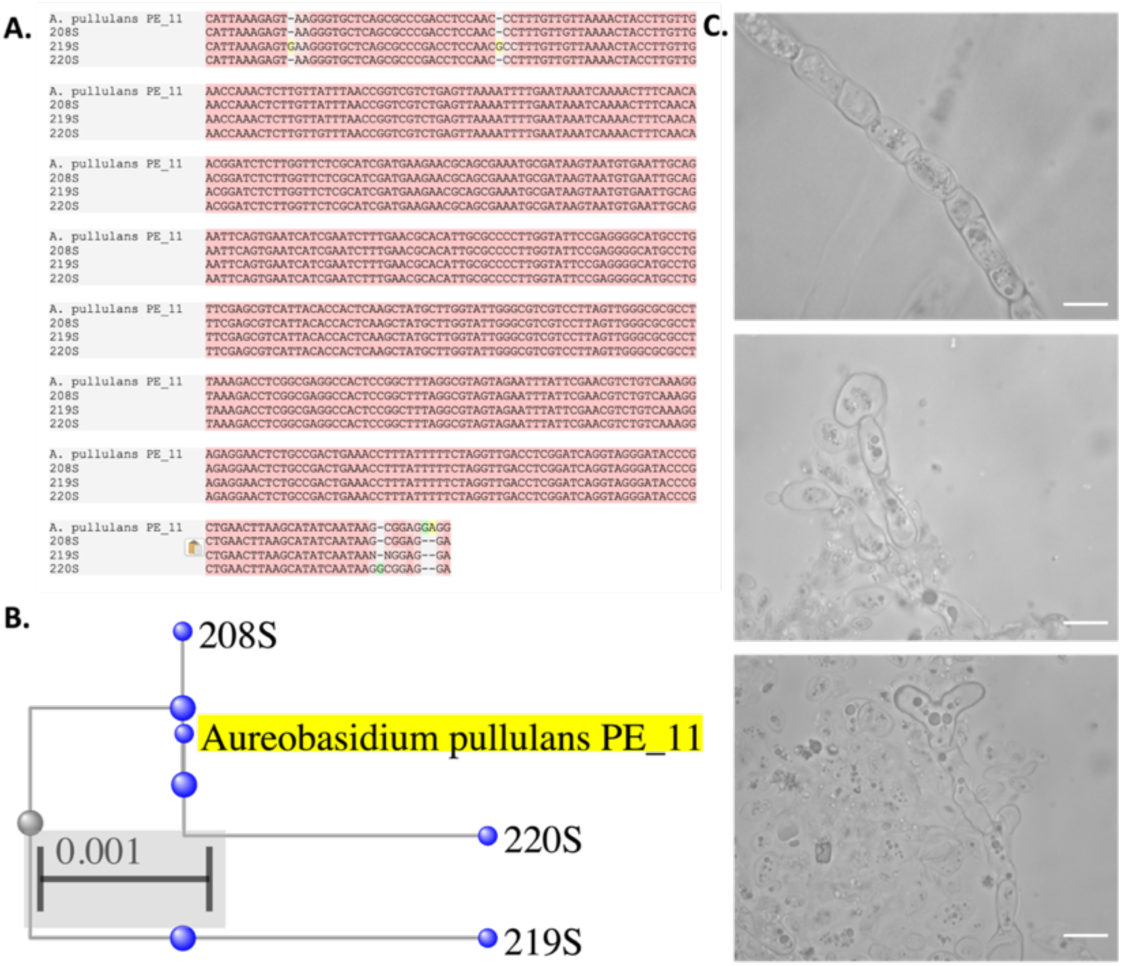
Identification of Aureobasidium pullulans from High-Temperature Environment. Alignment of three of the 15 A. pullulans cultures isolated from Fayette St. sidewalk (208S, 219S, 220S) to the A. pullulans PE_11 isolate (A), and their phylogenetic relationship (B). Microscopic images of the *A. pullulans* cultures show hyphal, yeast-like, and irregular shaped (C). Scale bar represents 20 µm.

